# LambdaPP: Fast and accessible protein-specific phenotype predictions

**DOI:** 10.1101/2022.08.04.502750

**Authors:** Tobias Olenyi, Céline Marquet, Michael Heinzinger, Benjamin Kröger, Tiha Nikolova, Michael Bernhofer, Philip Sändig, Konstantin Schütze, Maria Littmann, Milot Mirdita, Martin Steinegger, Christian Dallago, Burkhard Rost

**Author notes:** Correspondence with main authors through, http://www.rostlab.org/Tel: (email Rost). Equal first authorship (Tobias Olenyi, Céline Marquet). Equal senior authorship (Christian Dallago, Burkhard Rost).

## Abstract

The availability of accurate and fast Artificial Intelligence (AI) solutions predicting aspects of proteins are revolutionizing experimental and computational molecular biology. The webserver *LambdaPP* aspires to supersede PredictProtein, the first internet server making AI protein predictions available in 1992. Given a protein sequence as input, *LambdaPP* provides easily accessible visualizations of protein 3D structure, along with predictions at the protein level (GeneOntology, subcellular location), and the residue level (binding to metal ions, small molecules, and nucleotides; conservation; intrinsic disorder; secondary structure; alpha-helical and beta-barrel transmembrane segments; signal-peptides; variant effect) in seconds. The structure prediction provided by *LambdaPP* - leveraging *ColabFold and computed in minutes* - is based on *MMseqs2* multiple sequence alignments. All other feature prediction methods are based on the pLM *ProtT5*. Queried by a protein sequence, *LambdaPP* computes protein and residue predictions almost instantly for various phenotypes, including 3D structure and aspects of protein function.

**Accessibility Statement:** LambdaPP is freely available for everyone to use under embed.predictprotein.org, the interactive results for the case study can be found under https://embed.predictprotein.org/o/Q9NZC2. The frontend of LambdaPP can be found on GitHub (github.com/sacdallago/embed.predictprotein.org), and can be freely used and distributed under the academic free use license (AFL-2). For high-throughput applications, all methods can be executed locally via the bio-embeddings (bioembeddings.com) python package, or docker image at ghcr.io/bioembeddings/bio_embeddings, which also includes the backend of LambdaPP.

**Impact Statement:** We introduce LambdaPP, a webserver integrating fast and accurate sequence-only protein feature predictions based on embeddings from protein Language Models (pLMs) available in seconds along with high-quality protein structure predictions. The intuitive interface invites experts and novices to benefit from the latest machine learning tools. LambdaPP’s unique combination of predicted features may help in formulating hypotheses for experiments and as input to bioinformatics pipelines.

## Introduction

### PP protein prediction since dawn of internet

Launched 30 years ago, the PredictProtein (PP) web server provides a comprehensive interface for protein sequence analysis (56; 59; 76; 12). As the first internet server for predicting aspects of protein structure and function, it offers a broad overview of predicted features. Amongst many innovations, PP introduced the combination of evolutionary information (EI) from multiple sequence alignments (MSAs) and machine learning (57), a subset of artificial intelligence (AI), for protein prediction. Nature’s 2021 method of the year (44), *AlphaFold2* (31), peaked the innovation by essentially solving the protein structure prediction problem with models approaching experimental high-resolution, inspiring a new era of advancing methods (2; 6; 46) and their application (17; 33; 78). *AlphaFold2* came when more sequences than ever before (2.1 billion proteins in BFD (67)) met new AI-optimized algorithms and hardware. *PredictProtein* and *AlphaFold2* work great in their domains and integrating both might help in enabling experts and novices alike to experiment, hypothesize, and generate novel insights quickly.

### EI+AI top, but not without caveats

Since the release of PredictProtein, the amount of non-annotated sequences has been rapidly increasing (58; 67). In fact, the sequence-annotation gap continues to grow despite experimental advances, e.g., experimental residue binding annotations are currently added for only two sequence-unique proteins per month for any organism and any ligand (40). AI models mitigate this gap. Until 2020, almost all state-of-the-art prediction methods had implemented the concept introduced by PP, namely inputting MSAs into AI. Although superfast tools relying on algorithmic and hardware advances sped up MSA generation (47; 16), biodatabases continue outgrowing the pace at which computer hardware accelerates (48; 71; 66). This challenge cannot be resolved by advancing computers. On top, MSAs are not always informative, especially for small sequence families, or proteins of the Dark Proteome (51).

### Protein Language Models (pLMs) solving problems?

Developments in representation learning (8), particularly in natural language processing (18), let to encoding latent protein information including aspects of evolutionary information. Protein language models (pLMs) based on deep learning large sets of unannotated sequences to generate numerical representations (embeddings) (9; 25; 41; 24; 50; 55). Embeddings from pLMs have been successfully used as input to downstream protein prediction tools (79; 38-40; 42; 43; 45; 65; 14; 26; 28; 64; 73). Some pLM-based methods still appear inferior to top MSA-based methods (24; 39; 73), others bested those (5; 10; 24; 40; 43; 65; 29; 30; 37). Performance has risen so much that sequence-specific pLM-based predictions now can capture some aspects of structural and functional dynamics better than much more accurate family-averaged solutions even from *AlphaFold2* (36; 73; 75).

### pLM-based protein predictions for the web

30 years ago, PredictProtein offered the first access to a variety of MSA-based AI solutions. Similarly, LambdaPP now makes state-of-the-art solutions for embedding-based predictions available. The server outputs predictions for the entire query protein (per-protein) and for each of its residues (per-residue, Fig. 1). All results are linked to 3D structure visualizations, currently retrieved from the AlphaFold Database (release 4, 07/2022 created using *AlphaFold Monomer v2*.*0 pipeline*) (72) or if unavailable predicted using *ColabFold* (v2.1.14) simplifying *AlphaFold2* at similar performance (46). Currently the only non-pLM method in the LambdaPP frame, it will soon be complemented by pLM-based solutions (36; 73; 75). As novel AI tools leveraging embeddings emerge, e.g. predicting CATH (63) classes (26), LambdaPP will be updated to extend its breadth. All feature prediction methods integrated into the LambdaPP webserver currently use ProtT5 (24) that, in our hands, outperformed ESM-1b (55) and others (4; 9; 25; 24) for numerous applications (38-40; 43; 65; 13; 26; 73). This consistency also increases speed as the generation of embeddings becomes a limiting step.

**Fig. 1:**
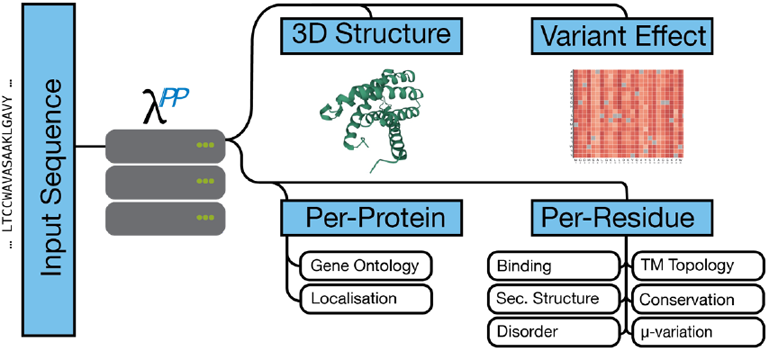
*LambdaPP* pipeline. Starting with an amino acid sequence, LambdaPP orchestrates the prediction of (1) protein 3D structure by ColabFold (46); (2) per-protein features: Gene Ontology (GO) annotations: goPredSim (39), subcellular location: LA (65); (3) per-residue features: binding residues: bindEmbed21DL (40), conservation: ProtT5cons (43), disorder: SETH (30), secondary structure: ProtT5-sec (24), helical and barrel transmembrane (TM) regions: TMbed (13); and (4) variant effect scores: VESPAl (43).

## Results

### Access of server

All methods are available through embed.predictprotein.org where users submit amino acid sequences up to 2000 residues. This limit speeds-up response-time (embedding computation is non-linear in protein length). Formats currently handled are: FASTA sequence, UniProt accession number, UniProt protein name (70), or a string of residues (“AA format”). Results are displayed immediately if cached, if not they are computed on the fly within seconds for pLM-based methods. There is no queuing system, as it takes longer to generate the display items on the frontend than to compute the predictions in the backend. Protein 3D structures are fetched from the AlphaFold Database (72) when inputting UniProt accessions, or predicted by ColabFold (46) through a *first come, first serve* queue (completing within 30 minutes for protein with 350 residues). Due to limited GPUs, *ColabFold* predictions are restricted to proteins shorter than 500 residues, but work is underway to transition to pLM-based 3D structure prediction providing fast and accurate predictions for longer sequences. All results are cached for 10 days before being deleted to conserve disk space and respect data privacy. Users can download results.

### Frontend and interface

The main LambdaPP interface displays the predictions in thematically ordered sections. Leveraging the *neXtProt* feature viewer (60), per-residue predictions are displayed in one view-pane (Fig. 2). The neXtProt plugin enables to display categorial features, e.g., binding, transmembrane-regions, and secondary structure as colored regions, and continues features, e.g., disorder, variance effect, and conservation, as line plots. An interactive connection between residue-level features and 3D structure maps predictions onto 3D visualization while displaying additional information in tooltips. The *protein-level section* visualizes subcellular location through colored images (20; 65) and GO-term predictions as lists of predicted GO-terms along with scores reflecting reliability (RI) and links to the reference protein used for the annotation transfer (39). The *Single Amino acid Variant (SAV) effect section* features the predictions of how much point mutations (SAVs) negatively affect molecular function. By default, the effect is predicted for all 19 non-native SAVs, i.e., all point mutants irrespectively of their reachability through single nucleic variants (SNVs or SNPs). Finally, the predicted 3D structure is visualized in the *structure section*, using the Mol* plugin (62). To facilitate exploring predictions, we offer two alternative interfaces: the *print-page*, which displays several residue-level features in a print-ready form, and the *interactive* page, which displays the neXtProt feature viewer and 3D structure prediction in a single panel to allow easier interactive exploration. These alternative displays are reached from the main interface by clicking on the suggested alternative display buttons.

**Fig. 2:**
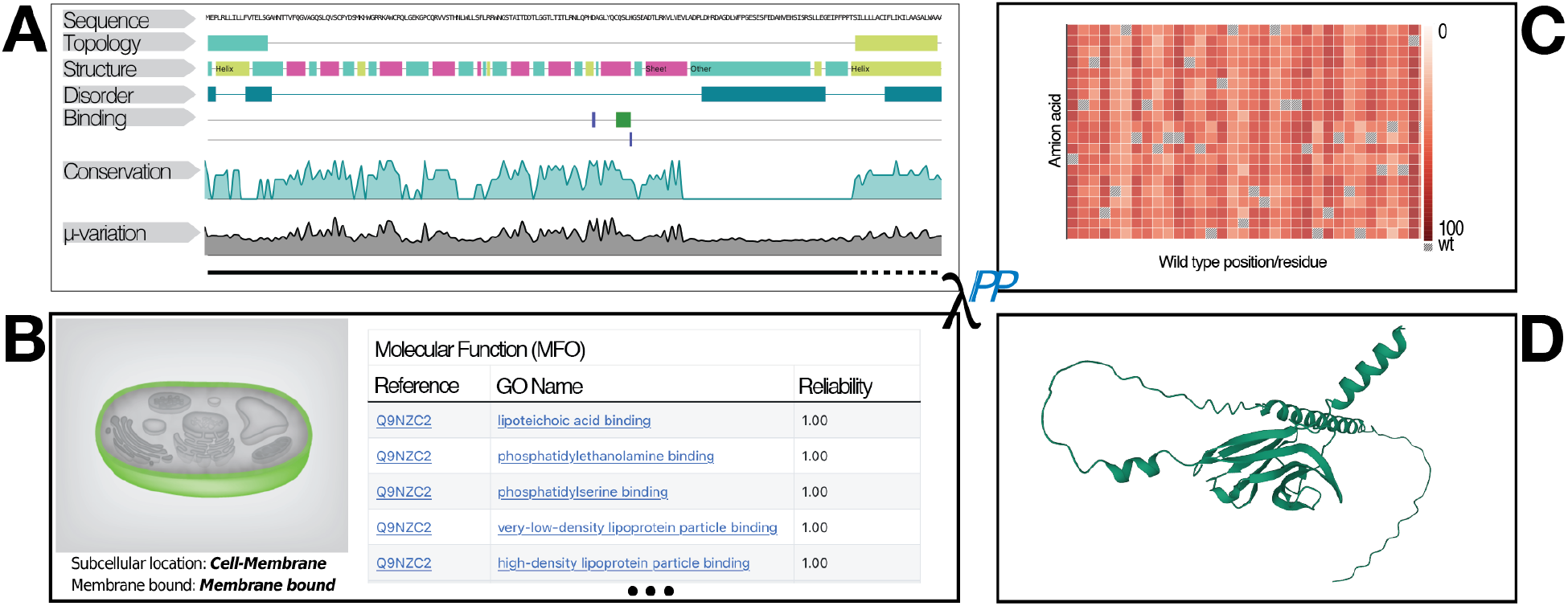
LambdaPP output for *TREM2_HUMAN*. *<u>Panel A:</u>* residue level features: secondary structure, transmembrane topology, disordered residues, small molecule, nucleic or metal binding residues, residue conservation and average variation (24; 40; 43; 13; 30); *<u>Panel B:</u>* sequence-level features: predicted subcellular localization (65), and an excerpt of predicted GO-annotations (39); *<u>Panel C:</u>* effect of SAVs (wild-type sequence on x-axis, mutations on y-axis; darker color=higher effect) (43); and *<u>Panel D</u>*: predicted 3D structure (46). Interactive version at https://embed.predictprotein.org/o/Q9NZC2.

### Backend and programmatic access

Users can retrieve prediction results programmatically through the bio-embeddings REST API (details: api.bioembeddings.com/api). This interface allows, e.g., to create ProtT5 embeddings of proteins with up to 2,000 residues (hardware not model restriction), and to retrieve the full spectrum of available annotations as generated by the backend, i.e., to download all predictions in JSON format. The backend could be hosted entirely on a standard workstation equipped with a workstation GPU (e.g., Quadro RTX 8000 46GB RAM), delivering results in seconds compared to minutes or hours for multi-node and cluster-based PP (12). However, to counter machine faults and guarantee availability, the hosted LambdaPP runs on different servers at the LCSB in Luxembourg and the TUM in Munich and can process one request at a time, on average, in four seconds for proteins of 350 residues. The hosted backend can be manually scaled to respond to parallel requests during high demand.

### Availability for local deployment

To take advantage of the methods feature on LambdaPP locally, advanced users can rely on the bio-embeddings package (21) (bioembeddings.com). Along with various use cases, it provides a docker image (ghcr.io/bioembeddings/bio_embeddings) for easy deployment on local machines. We recommend installing bio-embeddings locally to avoid length restrictions imposed on LambdaPP.

### Use case: *triggering receptor* (Q9NZC2)

We demonstrated the LambdaPP workflow using the *triggering receptor expressed on myeloid cells 2* (UniProt accession: Q9NZC2) protein and compared results to the expert curated UniProtKB entry (70). We selected Q9NZC2, as it is associated with *Polycystic lipomembranous osteodysplasia with sclerosing leukoencephalo-pathy* (PLOSL2) and has diverse annotations in different regions (all at SOM Fig. S1).

#### Per-protein

LambdaPP trivially listed most UniProtKB GO annotations with high reliability because the protein was in goPredSim’s lookup set. Subcellular location was correctly predicted as cell-membrane, and the structure from the AlphaFold Database (72) aligned well with the structures of 5ELI (3; 32) (RMSD: 0.45 Å) and 5UD8 (68; 69) (RMSD: 0.37 Å; SOM: Fig. S4).

#### Per-residue

LambdaPP marked the first 17 residues as signal peptides, matching the automatic (rule-based) UniProtKB annotation (one residue shorter). The transmembrane stretch matched with the UniProtKB transmembrane stretch (4 residues shorter; SOM: Fig. S5).

#### Binding

UniProtKB has no annotation of metal ion, small molecule, or nucleic acid binding, while bindEmbed21DL predicted two metal ion and one small-molecule binding residues with high reliability which might be interesting targets for future experiments (40).

#### Disorder & conservation

UniProtKB also annotates no intrinsic disorder. Yet, the predicted high disorder content for loop regions next to the transmembrane segment and the high order content outside correlated well with *AlphaFold 2’s* predicted *Local Distance Difference Test* (pLDDT), reflecting the confidence for the 3D prediction with pLDDT>70 typically considered reliable; Fig. S6) (52; 74). For sequence conservation, we compared the predictions shown by LambdaPP to those from ConSurfDB (7) obtained for 5ELI (3; 32) (mean squared error (MSE) ∼9) and 5UD8 (68; 69) (MSE ∼4; Fig. S7).

#### SAV effects

The predictions of the effects of single SAVs upon molecular function showed a similar trend as the UniProtKB annotations: Q9NZC2 seems susceptible to mutation effects. Zooming into residues relevant for binding to PLOS2, e.g., a mutation at residue position 126 (V>G) suggests a strong mutation effect (score 71). Residues marked in LambdaPP with high scores could be an interesting target for future mutational assays (e.g., residues at position 35, 85, 105).

### Interactive selection

When users select predicted residue features on the nextProt viewer, the selection is transferred into the 3D viewer (SOM: Fig. S3). This eases the identification of relevant structural regions. For instance, selecting the predicted signal peptide for Q9NZC2 (SOM: Fig. S3), highlights the region on 3D structure and allows to visually verify the prediction.

### Use case: predicting a long protein

We selected a hypothetical *ice nucleation* protein from *Pseudomonas syringae* (ICEV_PSESX, O33479) due to its length, its remarkable *AlphaFold2* structure prediction (Fig. 3A), and its possibly interesting ice-binding properties (19). Most of the structure is predicted with very high confidence (pLDDT>90; confirmed by predicted low disorder). Another region with low *AlphaFold2* pLDDT correlates with predicted low conservation and high disorder (residues 111-165, Fig. 3A: loop next to top left). Three residues are predicted as metal binding (41, 208, 1196), and thirteen as small molecule binding (173-175, 189-192, 222, 238-239, 253-254, 590, 606).

**Fig. 3:**
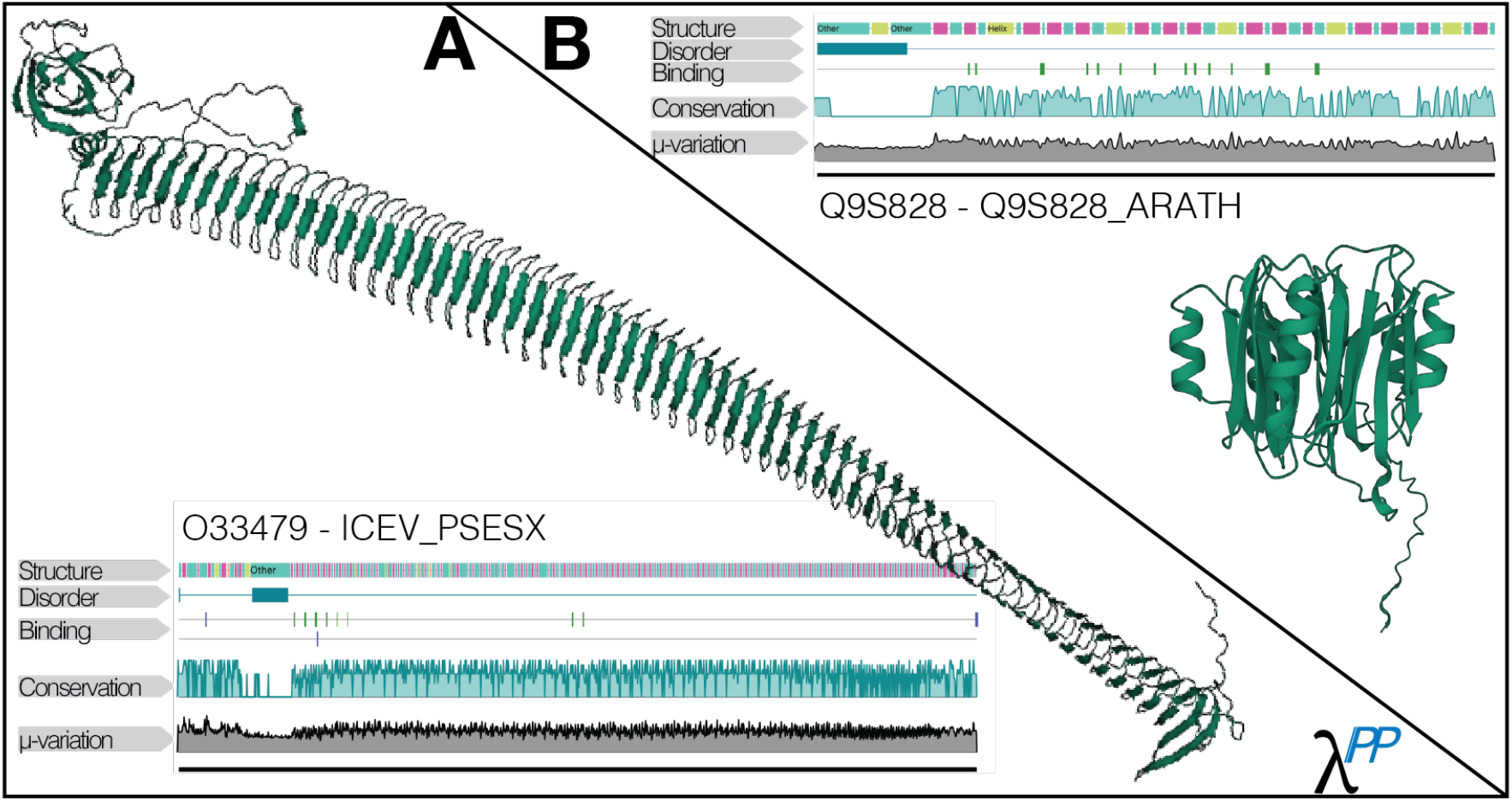
Remarkable AlphaFold2 predictions. *<u>Panel A</u>* (lower left triangle) displays the 3D structure predicted by AlphaFold2 for the ice nucleation protein ICEV_PSESX. The protein contains 1165 residues and is available through LambdaPP as part of AFDB. *<u>Panel B</u>* (upper right triangle) showcases the AlphaFold2 prediction of what might constitute a novel superfamily for the plant protein with the UniProt identifier Q9S828_ARATH.

#### Disaccord

*ProtT5Sec* predicted secondary structure differed partially from AlphaFold2 either suggesting alternative conformations or prediction inconsistencies. Similarly, GO annotations (CC: nucleus, BP: regulation of transcription by RNA polymerase II, MF: double-stranded DNA binding) are predicted with low reliability (RI: 0.24) and differ from those inferred by homology in UniProtKB (CC: cell outer membrane, MF: ice-binding). While low reliability is a good indicator that the predicted features should be taken with a grain of salt, the result remains interesting given the lack of proteins with reliable GO annotations for GOPredSim. This could point to understudied biological functions.

### Use case: annotating family and function for an unknown protein

A plant protein with UniProt ID Q9S828 remains uncharacterized by experiment. AlphaFold2’s 3D prediction suggests it folds into a new superfamily (15). The predicted pLDDT of the 3D structure (very low 1-29, low 30-33, very low 34-45, low 46-48) matches partially with UniProtKB disorder annotations (1-29) and LambdaPP-included disorder predictions (residue 1-39; N-terminal region looping to the lower left (Fig. 3B). This region is also predicted as poorly conserved.

LambdaPP suggests that Q9S828 might be a serine-threonine kinase (GO:0004674), involved in the phosphorylation of peptidyl-serine (GO:0018105) and contribute to flower development (GO:0009908). Compared to the broader UniprotKB annotations (GO:001630, GO:0016310), functional annotations provided by LambdaPP allow to hypothesize about the protein’s functional role in the plant and design targeted experiments to validate the predictions.

## Conclusions

LambdaPP importantly advances into the next generation of protein prediction, providing the first lightning fast, all-round protein prediction server based almost exclusively on embeddings from one protein language model – ProtT5, accompanied by high-quality 3D structure predictions from the *AlphaFold DB* or *ColabFold*. The web interface allows detailed analyses without requiring users to chip-in high-end servers, AI knowledge, or programming skills. The sub-minute turnaround time from sequence input to predictions of 15 different per-residue and per-protein features coupled with an intuitive user interface allow users to quickly generate an overview for any desired protein sequence, in turn allowing hypothesis generation and analysis. Thereby, LambdaPP might become valuable for bridging the sequence-annotation gap through predictions from high-quality methods, allowing researchers to prioritize experiments and curation efforts.

## Materials & Methods

LambdaPP computes protein language model (pLM) representations (embeddings) using ProtT5 (24) from single amino acid sequences. More specifically, the embeddings are derived exclusively from the encoder-part of ProtT5 in half-precision, i.e., from float32 to float16 model weights, speeding up inference and improving performance of subsequent models (24). These embeddings are input to all methods provided via LambdaPP except ColabFold (46). Per-protein features predicted solely with ProtT5 embeddings as input currently include subcellular location (65), and Gene Ontology terms (GO) (39). Per-residue predictions solely with ProtT5 embeddings as input include: conservation (43); helical transmembrane regions, transmembrane beta barrels, along with signal peptides (13); binding for various ligands (40); intrinsically disordered regions (30); secondary structure (24). LambdaPP also predicts the effect of introducing single amino acid variants (SAV) in the input sequence upon molecular function, which uses the predicted conservation with a BLOSUM62-score (27) of the SAV as input (43). Additionally, the 3D structure for the query-sequence is currently predicted with ColabFold (46) implementing AlphaFold2 (31), however pLM-based alternatives are in development and will be available online soon (73).

### Per-protein: Gene Ontology (GO)

The method goPredSim (39) predicts GO terms by transferring annotations from the closest neighbor in a lookup dataset of proteins with known GO annotations (39). The closest neighbor is defined by the smallest pairwise Euclidean distance calculated between the ProtT5 embeddings of the lookup set and the target. The distance is converted to a Reliability Index (RI) ranging from 0 (weak prediction) to 1 (confident prediction). RI values above 0.35 for biological process ontology (BPO), 0.28 for molecular function ontology (MFO), and 0.29 cellular component ontology (CCO) suggest reliable results. Replicating CAFA3 (80), goPredSim reached F_max_(BPO)=38±2%, MFO: 52±3%, and CCO: 59±2% using ProtT5 embeddings (12; 39). Tested on proteins annotated after Feb. 2020 and confirmed by CAFA4 (23), results were slightly better (github.com/Rostlab/goPredSim-performance-assessment).

### Per-protein: subcellular location

For a given protein sequence, light attention (LA) predicts where in a cell a protein functions, i.e., its subcellular location or cellular compartment (65). Ten subcellular localization classes are differentiated as mapped in DeepLoc (5). The LA network architecture using ProtT5 embeddings as input significantly outperformed MSA-based state-of-the-art (SOTA) methods by about eight percentage points (Q10). For this task, ProtT5 embeddings (24) significantly outperformed all other pLM embeddings as input for the same architecture (4; 9; 25; 24; 55).

### Per-residue: ligand-binding

bindEmbed21DL (40), a two-layer convolutional neural network (CNN) exclusively inputting ProtT5 embeddings, predicts residues binding to metal ions, nucleic acids, or small molecules, distinguishing the three classes. The pLM-based method substantially outperformed its MSA-based predecessor (F1=47±2% vs. F1=34±2%) (61) on binding annotations from BioLiP (77). The seemingly low F1-score hid that the method often outperformed human annotations, in the sense that all strongly predicted (high reliability) residues annotated as non-binding investigated in detail appeared to reveal missing annotations rather than prediction mistakes.

### Per-residue: conservation

ProtT5cons (43) is a two-layer CNN, predicting the degree to which a residue is conserved in an MSA without using an MSA as input. The conservation level is scaled from 0 (highly variable) to 8 (highly conserved) similarly to ConSurf-DB (7). ProtT5 embeddings outperformed those from the pLM ProtBERT and performed *on par* with the ESM-1b (55) pLM embeddings. While only taking embeddings as input, the performance of ProtT5cons was similar to ConSeq (11) using MSAs (two-state Matthews Correlation Coefficient MCC(embeddings)= 0.596±0.006 vs. MCC(ConSeq) = 0.608±0.006) when compared to conservation levels of ConSurf-DB.

### Per-residue: intrinsic disorder

SETH (30), a two-layer CNN, predicts the degree of intrinsic disorder of a residue as defined by the chemical shift Z-scores (CheZOD) (49), where values below 8 signify disorder and values above 8 signify order. Different pLMs were compared (ProtT5 (24), ProSE(10), ESM-1b (55), ProtBERT (24), SeqVec (25)) with ProtT5 numerically outperforming the others. SETH outperformed all existing SOTA approaches in terms of mean AUC (area under the receiver operating characteristic curve) and Spearman correlation (0.72±0.01 for SETH vs. 0.67±0.01 for next best method ODinPred (22)) as well as similar current solutions operating on ESM-1b embeddings (53).

### Per-residue: secondary structure

ProtT5-sec(24), a two-layer CNN, reached a Q3 (three state per-residue accuracy) of 81±1.6% for the CASP12 (1) test set and Q3 of 84±0.5% for a larger data set NEW364 (24) competitive with, or even surpassing, top methods relying on MSAs.

### Per-residue: transmembrane helices and strands

TMbed predicts for each residue one of four classes: alpha helical transmembrane (TM) region, transmembrane beta strand, signal peptide, or other (13). For proteins with TM regions, it also predicts the inside/outside orientation within the membrane, i.e., on which side of the membrane the N-terminus begins. The model uses a four-layer CNN combined with a Gaussian filter and a Viterbi decoder. When applied to a non-redundant test set, TMbed correctly predicted 94±8% of beta-barrel transmembrane proteins (TMPs) and 98±1% of alpha-helical TMPs at false positive rates <1%. Furthermore, TMbed placed on average 9 out of 10 transmembrane segments within five residues of the experimental observation. TMbed performed *on par* with or better than SOTA methods. It stood out in terms of its low false positive rate and speed; both making TMbed well suited for high-thruput annotation and filtering, such as annotating millions of AlphaFold2 models.

### Per-residue: variant effect and μ-variation

To predict the effect of SAVs, VESPAl (43) takes the nine-state conservation prediction by ProtT5cons, and the BLOSUM62 substitution matrix (27) as input for a logistic regression. The simple architecture of VESPAl was close in performance to SOTA MSA-based methods (54; 35), and the embedding-based ESM-1v (45) on 39 deep mutational scanning (DMS) experiments (with 135,665 SAV) that had not been used for development. The per-residue μ-variation describes the average effect score of the 19 possible substitutions for the respective wild type.

### *3D* structure prediction

LambdaPP also includes SOTA MSA-based 3D structure predictions. If UniProt accessions are used as an input, the 3D structure is retrieved from the AlphaFold Database (AFDB) (72). If the structure is unavailable in AFDB or the input is not a UniProt accession number, ColabFold (46) is employed to predict the 3D structure, easing access to AlphaFold2 (31). Toward this end, MSAs are generated by searching with MMseqs2 (47) against UniRef 30 and ColabFoldDB (70). For single predictions, this combination is 20-30 times faster than the original AlphaFold2 at little loss of performance on the CASP14 (34) targets (47). Further parameters are an early stop criterion of a prediction certainty (pLDDT) above 85 or below 40, a default recycle count of 3 and a compilation of only the best performing out of five AlphaFold2 models. As the goal of LambdaPP is to provide a single reference for pLM-based predictions, 3D structure will soon be predicted using tools just recently presented in the literature (36; 73; 75), which will allow structure prediction to happen in seconds rather than in minutes, at accuracy comparable with MSA-based methods.

## Supporting information

SOM Figures S1,2,3,5

SOM main

## Abbreviations used

3D: three-dimensional
AI: Artificial Intelligence
AlphaFold2: break-through method for protein 3D structure prediction (31)
API: application programming interface
AUC: area under the receiver operating characteristic curve
BPO: biological process ontology
CAFA: critical assessment of functional annotation
CASP: critical assessment of protein structure prediction
CATH: class, architecture, topology, homologous superfamily
CCO: cellular component ontology
CheZOD: chemical shift Z-scores
CI: confidence interval
CNN: convolutional neural network
ColabFold: protocol for fast execution of *AlphaFold2* (46)
DB: database
DMS: deep mutational scanning
EI: evolutionary information
LambdaPP: embedding based protein predictions
GO: gene ontology
GPU: graphical processing unit
LA: light attention
MFO: molecular function ontology
MSA: multiple sequence alignment
pLDDT: predicted Local Distance Difference Test
pLM: protein language model
PP: PredictProtein
RI: Reliability Index
RMSD: root-mean-square deviation
SAV: single amino variant effect
SE: standard error
SOTA: state-of-the-art
TM: transmembrane
TMP: transmembrane protein
TPU: tensor processing unit

## Supplementary Material (SOM)

The Supplementary Online Material (SOM) provides additional display items, SOM_figures.pdf contains the UniProtKB Annotations at submission (Figure S1), a full print-out of LambdaPP predictions for TREM2_HUMAN (Figure S2), an example for the structure exploration pane for TREM2_HUMAN (Figure S3), and aligned UniProtKB and LambdaPP annotations (Figure S5). Additional details are provided in the main document (SOM.pdf).

## Acknowledgements

Thanks primarily to Tim Karl (TUM) for invaluable help with hardware and software and to Inga Weise (TUM) for support with many other aspects of this work. Thanks to John Jumper and his team at DeepMind for the breakthrough development of AlphaFold2 and for making code and predictions freely available. Two anonymous reviews helped substantially: thanks! Last, not least, thanks to all who deposit experimental data in public databases, to those who maintain these databases, and those who make methods available that allow enriching the experimental data.

