## Supplementary material for "LambdaPP: Fast and accessible protein-specific phenotype predictions": SOM Figures S1,2,3,5

Lamdba Predict Protein (λPP)    Publication    Code

Using sequence with Accession Q9NZC2.

MEPLRLLILLFVTELSGAHNTTVFQGVAGQSLQVSCPYDSMKHWGRRKAWCRQLGEKGPCQRVVSTHNLWLLSFLRRWNGS
TAITDDTLGGTLTITLRNLQPHDAGLYQCQSLHGSEADTLRKVLVEVLADPLDHRDAGDLWFPGESESFEDAHVEHSISRSLLEG
EIPFPPTSILLLLACIFLIKILAASALWAAAWHGQKPGTHPPSELDCGHDPGYQLQTLPGLRDT

New Query

Overview    Interactive    Print

### Residue Features

Method Details

Download all predictions

**Position :** 0    **Zoom :** x 1

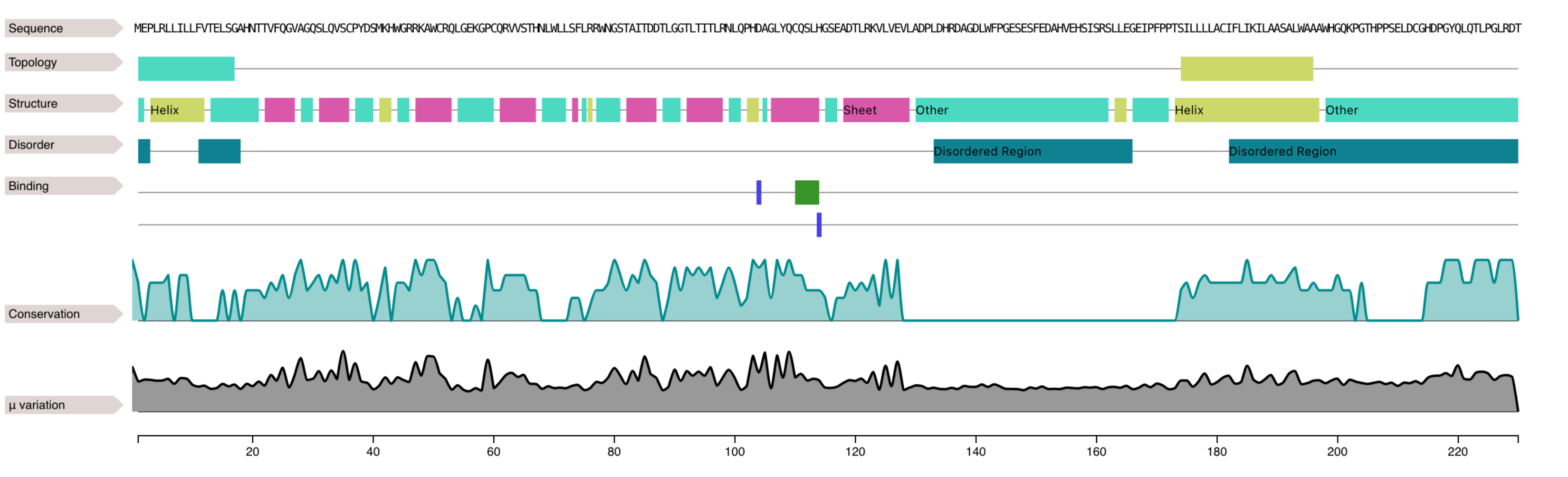


### Protein Features

Subceullular Location

Method Details

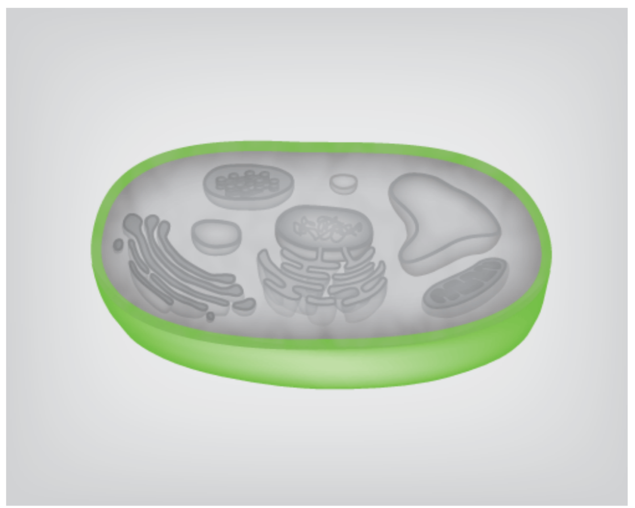

Subcellular location: **Cell-Membrane**

Membrane bound: **Membrane bound**

Gene Ontology

Method Details

**Biological process (BPO)**

| Reference Seq. | GO Name | Reliability Index |
|---|---|---|
| **Q9NZC2** | positive regulation of protein localization to plasma membrane | 1.00 |
| **Q9NZC2** | memory | 1.00 |
| **Q9NZC2** | response to axon injury | 1.00 |
| **Q9NZC2** | negative regulation of p38MAPK cascade | 1.00 |
| **Q9NZC2** | apoptotic cell clearance | 1.00 |
| **Q9NZC2** | negative regulation of macrophage colony-stimulating factor signaling pathway | 1.00 |
| **Q9NZC2** | positive regulation of ATP biosynthetic process | 1.00 |
| **Q9NZC2** | import into cell | 1.00 |
| **Q9NZC2** | cellular response to lipid | 1.00 |
| **Q9NZC2** | negative regulation of autophagy | 1.00 |
| **Q9NZC2** | cellular response to lipoteichoic acid | 1.00 |
| **Q9NZC2** | negative regulation of inflammatory response to antigenic stimulus | 1.00 |
| **Q9NZC2** | positive regulation of chemotaxis | 1.00 |
| **Q9NZC2** | positive regulation of TOR signaling | 1.00 |
| **Q9NZC2** | complement-mediated synapse pruning | 1.00 |
| **Q9NZC2** | negative regulation of amyloid fibril formation | 1.00 |
| **Q9NZC2** | neuroinflammatory response | 1.00 |
| **Q9NZC2** | regulation of macrophage inflammatory protein 1 alpha production | 1.00 |
| **Q9NZC2** | regulation of oxidative stress-induced neuron intrinsic apoptotic signaling pathway | 1.00 |
| **Q9NZC2** | negative regulation of autophagic cell death | 1.00 |
| **Q9NZC2** | humoral immune response | 1.00 |
| **Q9NZC2** | cellular response to glucose stimulus | 1.00 |
| **Q9NZC2** | positive regulation of complement activation, classical pathway | 1.00 |
| **Q9NZC2** | cellular response to amyloid-beta | 1.00 |
| **Q9NZC2** | astrocyte activation | 1.00 |
| **Q9NZC2** | negative regulation of cholesterol storage | 1.00 |
| **Q9NZC2** | positive regulation of inward rectifier potassium channel activity | 1.00 |
| **Q9NZC2** | detection of peptidoglycan | 1.00 |
| **Q9NZC2** | defense response to Gram-negative bacterium | 1.00 |
| **Q9NZC2** | positive regulation of osteoclast differentiation | 1.00 |
| **Q9NZC2** | negative regulation of toll-like receptor 4 signaling pathway | 1.00 |
| **Q9NZC2** | positive regulation of phosphatidylinositol 3-kinase signaling | 1.00 |
| **Q9NZC2** | negative regulation of tumor necrosis factor production | 1.00 |
| **Q9NZC2** | positive regulation of CD40 signaling pathway | 1.00 |
| **Q9NZC2** | positive regulation of cholesterol efflux | 1.00 |
| **Q9NZC2** | positive regulation of antigen processing and presentation of peptide antigen via MHC class II | 1.00 |
| **Q9NZC2** | positive regulation of microglial cell activation | 1.00 |
| **Q9NZC2** | detection of lipoteichoic acid | 1.00 |
| **Q9NZC2** | dendritic spine maintenance | 1.00 |
| **Q9NZC2** | respiratory burst after phagocytosis | 1.00 |
| **Q9NZC2** | osteoclast differentiation | 1.00 |
| **Q9NZC2** | positive regulation of engulfment of apoptotic cell | 1.00 |
| **Q9NZC2** | positive regulation of microglial cell migration | 1.00 |
| **Q9NZC2** | negative regulation of glial cell apoptotic process | 1.00 |
| **Q9NZC2** | positive regulation of CAMKK-AMPK signaling cascade | 1.00 |
| **Q9NZC2** | regulation of toll-like receptor 6 signaling pathway | 1.00 |
| **Q9NZC2** | phagocytosis, engulfment | 1.00 |
| **Q9NZC2** | microglial cell activation involved in immune response | 1.00 |
| **Q9NZC2** | regulation of lipid metabolic process | 1.00 |
| **Q9NZC2** | social behavior | 1.00 |
| **Q9NZC2** | positive regulation of peptidyl-tyrosine phosphorylation | 1.00 |
| **Q9NZC2** | regulation of plasma membrane bounded cell projection organization | 1.00 |
| **Q9NZC2** | dendritic cell differentiation | 1.00 |
| **Q9NZC2** | positive regulation of low-density lipoprotein particle clearance | 1.00 |
| **Q9NZC2** | positive regulation of high-density lipoprotein particle clearance | 1.00 |
| **Q9NZC2** | negative regulation of NLRP3 inflammasome complex assembly | 1.00 |
| **Q9NZC2** | positive regulation of macrophage fusion | 1.00 |
| **Q9NZC2** | regulation of resting membrane potential | 1.00 |
| **Q9NZC2** | negative regulation of astrocyte activation | 1.00 |
| **Q9NZC2** | negative regulation of toll-like receptor 2 signaling pathway | 1.00 |
| **Q9NZC2** | detection of lipopolysaccharide | 1.00 |
| **Q9NZC2** | pyroptosis | 1.00 |
| **Q9NZC2** | positive regulation of proteasomal protein catabolic process | 1.00 |
| **Q9NZC2** | negative regulation of sequestering of triglyceride | 1.00 |
| **Q9NZC2** | positive regulation of ERK1 and ERK2 cascade | 1.00 |
| **Q9NZC2** | CXCL12-activated CXCR4 signaling pathway | 1.00 |
| **Q9NZC2** | positive regulation of synapse pruning | 1.00 |
| **Q9NZC2** | positive regulation of amyloid-beta clearance | 1.00 |
| **Q9NZC2** | phagocytosis, recognition | 1.00 |
| **Q9NZC2** | negative regulation of interleukin-1 beta production | 1.00 |
| **Q9NZC2** | positive regulation of protein secretion | 1.00 |
| **Q9NZC2** | regulation of innate immune response | 1.00 |
| **Q9NZC2** | amyloid-beta clearance by cellular catabolic process | 1.00 |
| **Q9NZC2** | positive regulation of C-C chemokine receptor CCR7 signaling pathway | 1.00 |
| **Q9NZC2** | negative regulation of fat cell proliferation | 1.00 |
| **Q9NZC2** | negative regulation of I-kappaB kinase/NF-kappaB signaling | 1.00 |
| **Q9NZC2** | regulation of interleukin-6 production | 1.00 |
| **Q9NZC2** | positive regulation of interleukin-10 production | 1.00 |
| **Q9NZC2** | excitatory synapse pruning | 1.00 |
| **Q9NZC2** | response to ischemia | 1.00 |
| **Q9NZC2** | regulation of hippocampal neuron apoptotic process | 1.00 |
| **Q9NZC2** | negative regulation of phosphatidylinositol 3-kinase signaling | 1.00 |
| **Q9NZC2** | positive regulation of mitochondrion organization | 1.00 |
| **Q9NZC2** | lipid homeostasis | 1.00 |
| **Q9NZC2** | cellular response to oxidised low-density lipoprotein particle stimulus | 1.00 |
| **Q9NZC2** | positive regulation of kinase activity | 1.00 |
| **Q9NZC2** | cellular response to peptidoglycan | 1.00 |
| **Q9NZC2** | microglial cell proliferation | 1.00 |
| **Q9NZC2** | cellular response to hypoxia | 1.00 |
| **Q9NZC2** | negative regulation of cytokine production involved in inflammatory response | 1.00 |
| **Q9NZC2** | positive regulation of NIK/NF-kappaB signaling | 1.00 |

**Molecular function (MFO)**

| Reference Seq. | GO Name | Reliability Index |
|---|---|---|
| **Q9NZC2** | lipoteichoic acid binding | 1.00 |
| **Q9NZC2** | phosphatidylethanolamine binding | 1.00 |
| **Q9NZC2** | phosphatidylserine binding | 1.00 |
| **Q9NZC2** | very-low-density lipoprotein particle binding | 1.00 |
| **Q9NZC2** | high-density lipoprotein particle binding | 1.00 |
| **Q9NZC2** | scaffold protein binding | 1.00 |
| **Q9NZC2** | kinase activator activity | 1.00 |
| **Q9NZC2** | peptidoglycan binding | 1.00 |
| **Q9NZC2** | lipopolysaccharide binding | 1.00 |
| **Q9NZC2** | low-density lipoprotein particle binding | 1.00 |
| **Q9NZC2** | sulfatide binding | 1.00 |
| **Q9NZC2** | apolipoprotein A-I binding | 1.00 |
| **Q9NZC2** | transmembrane signaling receptor activity | 1.00 |
| **Q9NZC2** | amyloid-beta binding | 1.00 |
| **Q9NZC2** | protein tyrosine kinase binding | 1.00 |

**Cellular Component (CCO)**

| Reference Seq. | GO Name | Reliability Index |
|---|---|---|
| **Q9NZC2** | extracellular region | 1.00 |
| **Q9NZC2** | plasma membrane raft | 1.00 |
| **Q9NZC2** | integral component of plasma membrane | 1.00 |
| **Q9NZC2** | plasma membrane | 1.00 |

Single Amino Acid Variant Effect

Method Details

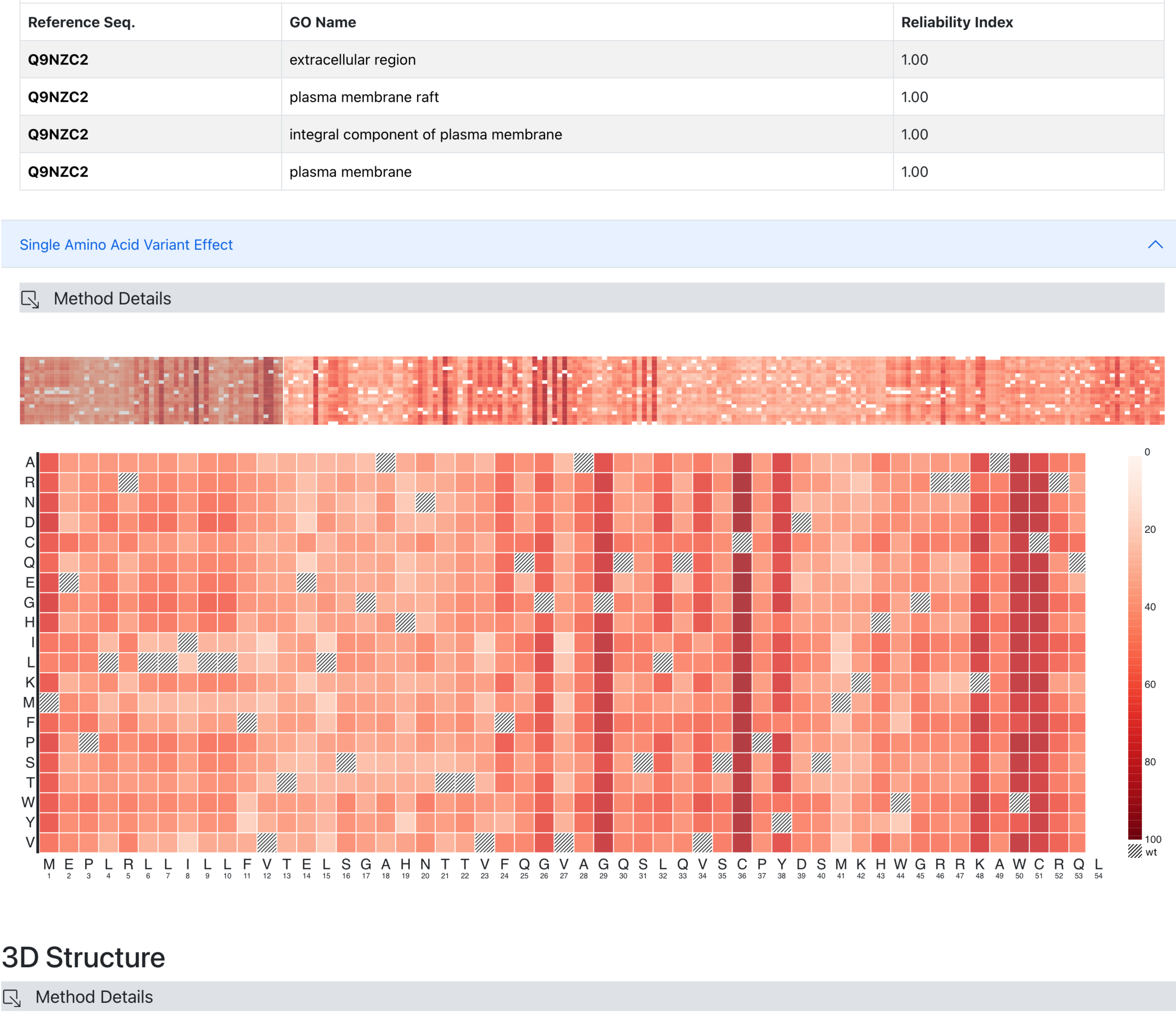


### 3D Structure

Method Details

Download Structure

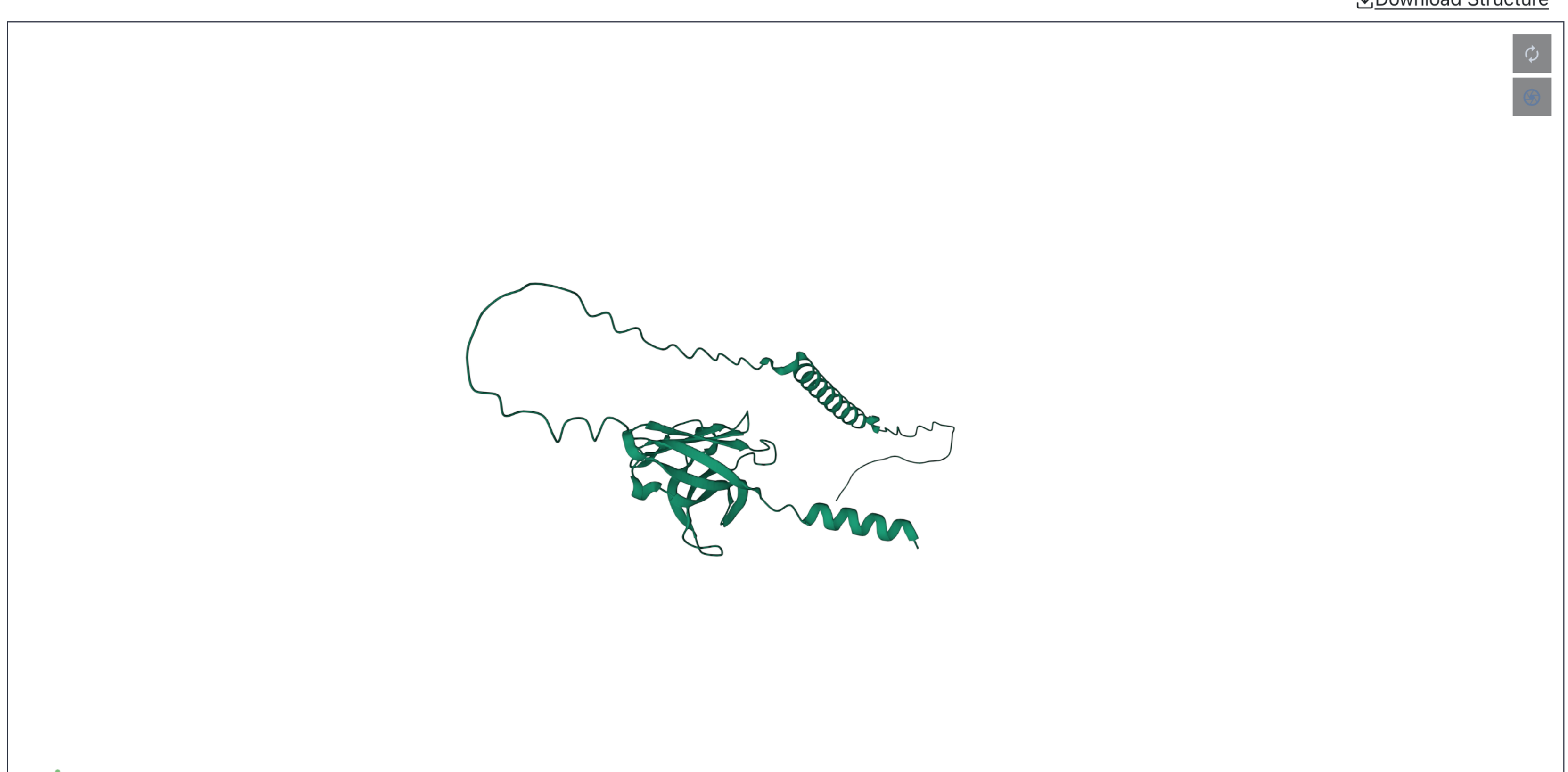

LINKS
Impressum
ToS & Privacy Policy
Contribute
Cite

SERVICES
Gloassary
Colab Fold
AlphaFold2 DB
UniprotKB

FOLLOW US

© 2022 Copyright: Rostlab

Using sequence with Accession **Q9NZC2**.

MEPLRLLILLFVTELSGAHNTTVFQGVAGQSLQVSCPYDSMKHWGRRKAWCRQLGEKGPCQRVVSTHNLWLLSFLRRWNGS
TAITDDTLGGTLTITLRNLQPHDAGLYQCQSLHGSEADTLRKVLVEVLADPLDHRDAGDLWFPGESESFEDAHVEHSISRSLLEG
EIPFPPTSILLLLACIFLIKILAASALWAAAWHGQKPGTHPPSELDCGHDPGYQLQTLPGLRDT

New Query

Overview Interactive Print

Explore predicted per-residue features on the structure by highlighting regions of interest in the feature viewer. Brush to zoom in, right click to zoom out.

Download all predictions

**Position :** 198W **Zoom :** x 1

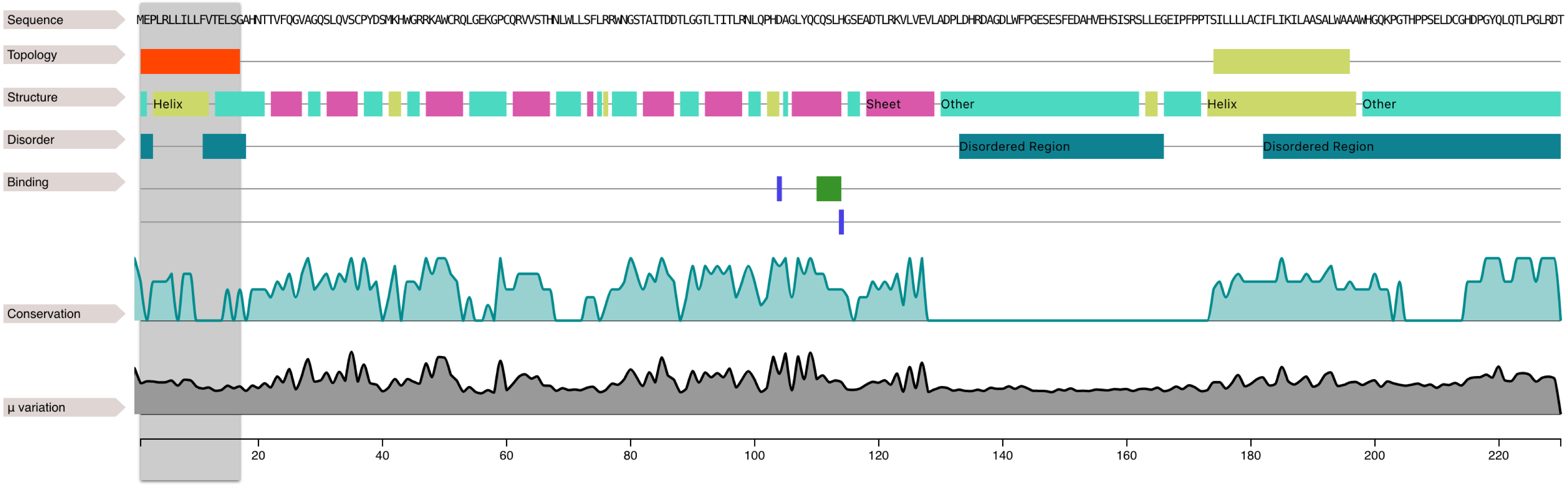


Download Structure

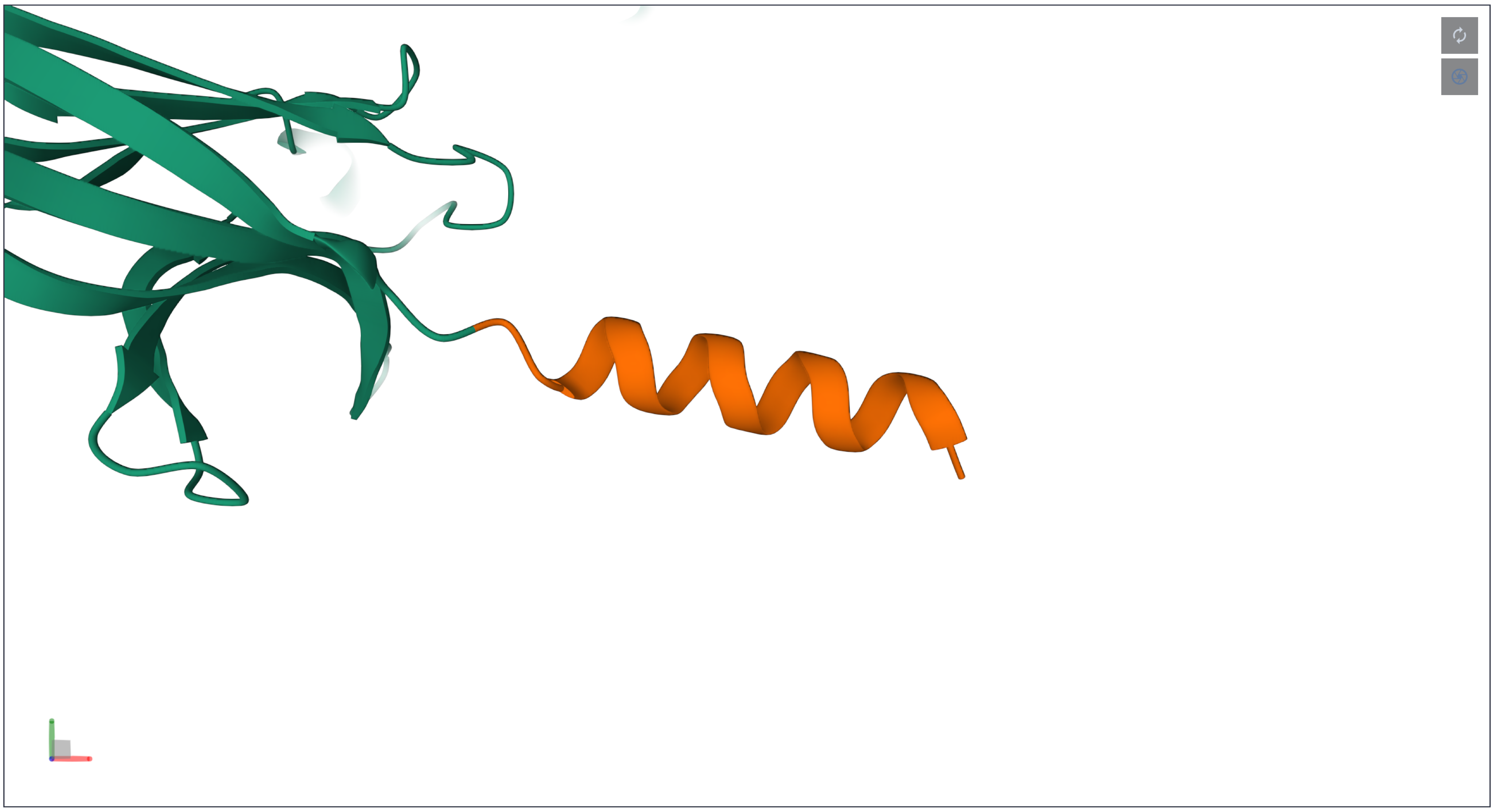



© 2022 Copyright: Rostlab

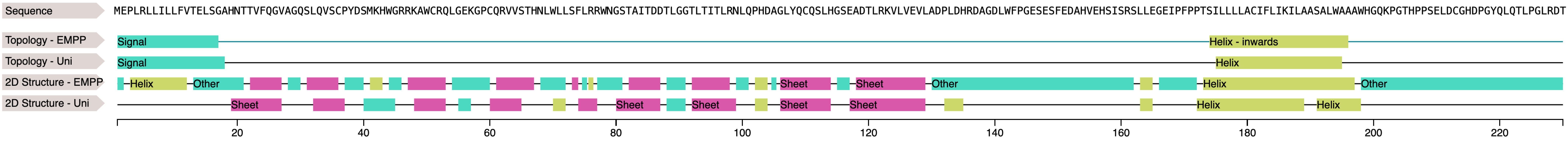
Sequence
MEPLRLLILLFVTELSGAHNTTVFQGVAGQSLQVSCPYDSMKHWGRRKAWCRQLGEKGPCQRVVSTHNLWLLSFLRRWNGSTAITDDTLGGTLTITLRNLQPHDAGLYQCQSLHGSEADTLRKVLVEVLADPLDHRDAGDLWFPGESESFEDAHVEHSISRSLLEGEIPFPPTSILLLLACIFLIKILAASALWAAAWHGQKPGTHPPSELDCGHDPGYQLQTLPGLRDT
Topology - EMPP
Signal
Helix - inwards
Topology - Uni
Signal
Helix
2D Structure - EMPP
Helix
Other
Sheet
Sheet
Other
Helix
Other
2D Structure - Uni
Sheet
Sheet
Sheet
Sheet
Sheet
Helix
Helix
20
40
60
80
100
120
140
160
180
200
220
