## Supplementary material for "LambdaPP: Fast and accessible protein-specific phenotype predictions": SOM main

### Supporting online material
### LambdaPP: Fast and accessible protein-specific phenotype predictions

Tobias Olenyi, Céline Marquet, Michael Heinzinger, Benjamin Kröger, Tiha Nikolova, Michael Bernhofer, Philip Sändig, Konstantin Schütze, Maria Littmann, Konstantin Weißenow, Milot Mirdita, Martin Steinegger, Christian Dallago & Burkhard Rost

#### Table of Contents for Supporting Online Material

#### Short description of Supporting Online Material

In this document, we provide figure captions for screenshots that were submitted as additional supporting file (SOM_figures.pdf; containing Fig. S1, S2, S3, S5) outside this document, and expand the case-study analyses with further details (Fig. S4, S6, S7). Finally, we give some background information on the structure prediction capabilities of LambdaPP.

### Material

#### 1.1 Caption for Fig. S1

**Fig. S1 (additional file): Screenshot of UniprotKB for TREM2_HUMAN (as of 31.7.2022).** Complete UniprotKB (10) entry for TREM2_HUMAN (Q9NZC2) with an annotation score of 5/5.

#### 1.2 Caption for Fig. S2

**Fig. S2 (additional file): Screenshot of LambdaPP for TREM2_HUMAN (as of 31.7.2022).** Complete Interface of LambdaPP on the example of TREM2_HUMAN (Q9NZC2). An interactive version of the displayed results can be found under https://embed.predictprotein.org/o/Q9NZC2 .

#### 1.3 Caption for Fig. S3

**Fig. S3 (additional file): Screenshot of LambdaPP for TREM2_HUMAN in interactive mode (as of 31.7.2022).** The structure exploration view of LambdaPP on the example of TREM2_HUMAN (Q9NZC2). The interactive mode allows to remove obtrusive information from the prediction page to show structure and feature view in proximity. This view enables quick exploration of individual features on the structure (and vice-versa) without scrolling.

#### 1.4 Structural Alignments between LambdaPP prediction and UniprotKB annotated structures

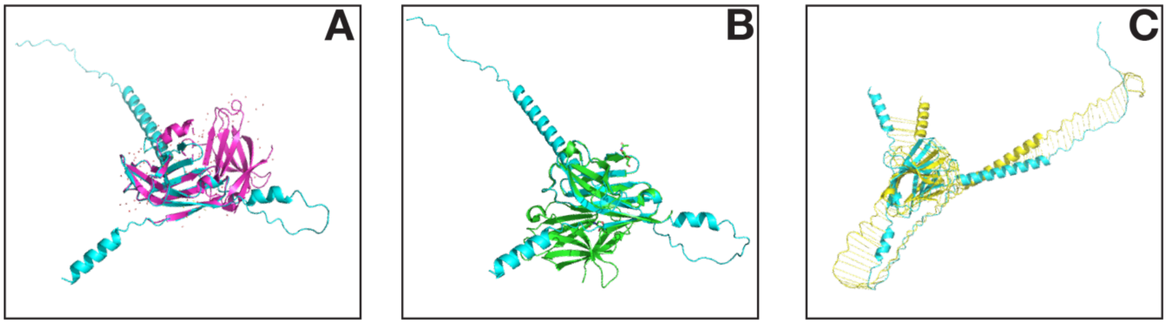

**Fig. S4: Structural alignment of LambdaPP prediction and structures annotated in UniprotKB.** *Panel A:* The structural alignment between the structure generated by LambdaPP (ColabFold (4)) and PDB-ID 5UD8 (8; 9) as computed by PyMol (6); *Panel B:* The alignment between the predicted structure and 5ELI (1; 3); *Panel C:* The alignment with the predicted AlphaFold 2 structure.

#### 1.5 Caption for Fig. S5

**Fig. S5 (additional file): Comparison of Predicted vs. Annotated Features**. The topology and structural annotations as predicted through LambdaPP and as annotated in UniprotKB aligned for convenient comparison. The annotations are visualized using Mol* (7). The information display highlights the close match of the predictions with the annotations in UniProtKB, especially for topology but also for secondary structure predictions.

#### 1.6 Comparison of Predicted Disorder with PlDDT as predicted by AlphaFold 2

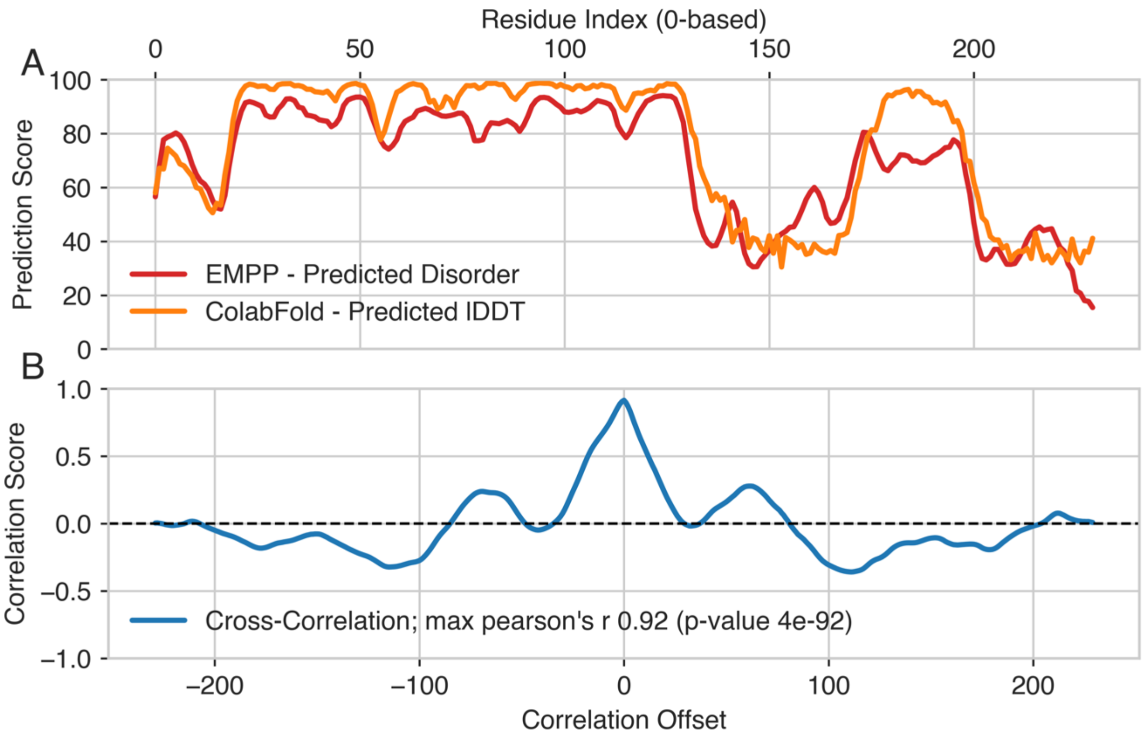

**Fig. S6: Comparison of Predicted Disorder with PlDDT as predicted by ColabFold.** *Panel A*: The predicted lDDT score of the structure prediction for TREM2_HUMAN (generated by ColabFold (4)) and the predicted disorder in EMPP normalized to a range between 0 and 100; *Panel B:* The normalized cross-correlation between the two signals. The high correlation between the two prediction signals indicates the accuracy of the disorder prediction (5; 12).

#### 1.7 Comparison of Predicted Conservation Scores with ConsurfDB

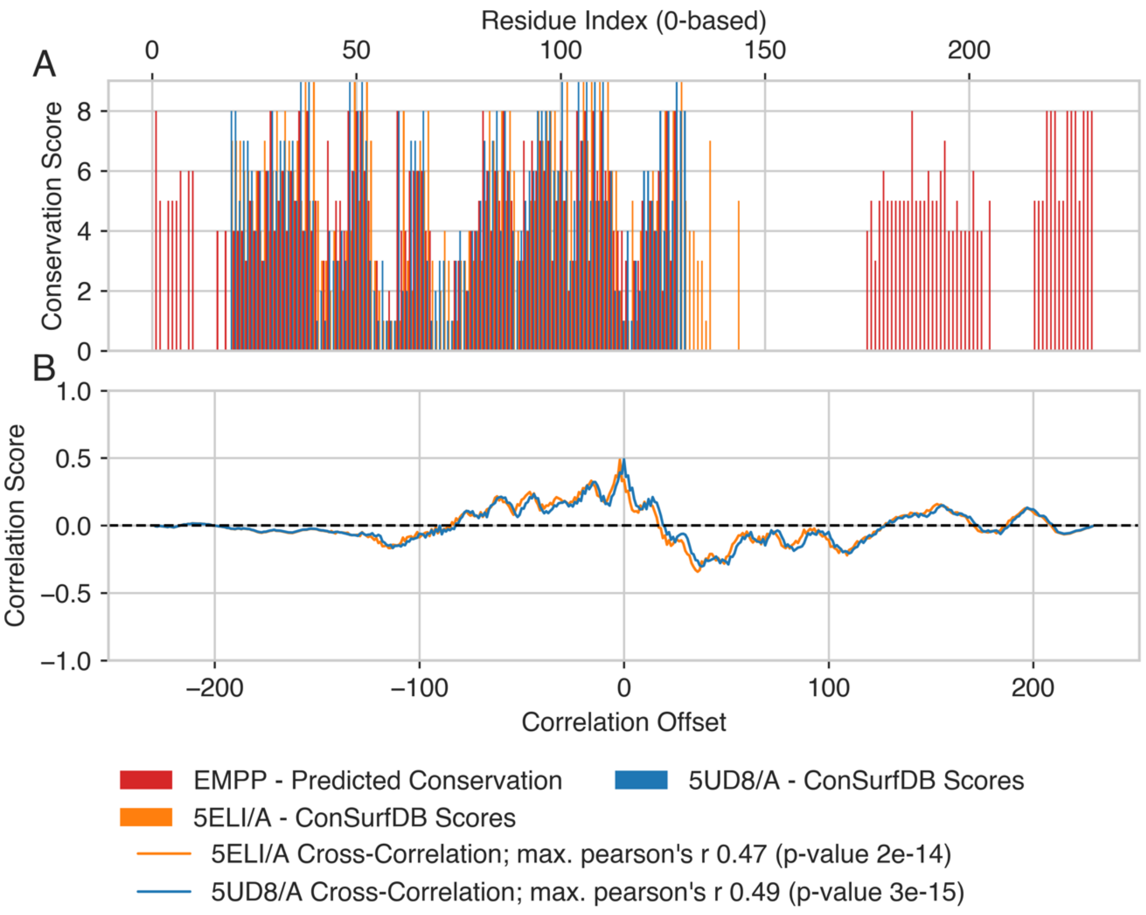

**Fig. S7: Comparison of Predicted Conservation Scores with ConSurfDB.** *Panel A:* The predicted conservation scores for TREM2_HUMAN, as well as the conservation scores for 5UD8 (8; 9) and 5ELI (1; 3) as determined by ConSurfDB (2); *Panel B:* The normalized cross-correlation between the predicted conservation and the ConSurfDB scores.

#### 1.8 Structure Retrieval / Prediction in LambdaPP

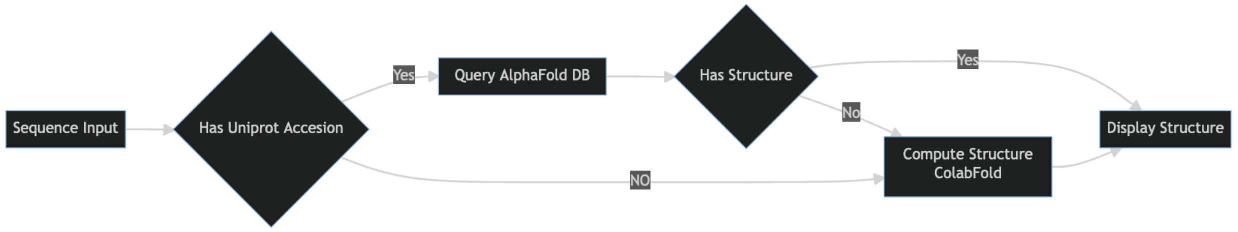

**Fig. S8: Workflow for structure Retrieval in LambdaPP.** If an accession is available, LambdaPP will first try to retrieve a matching structure from AlphaFold DB (11). If no structure can be found there, or no accession was available, LambdaPP will start a structure prediction Job based on ColabFold.

To display the predicted structure for a given sequence, LambdaPP queries AlphaFold DB (11), if possible. Should this not yield a viable structure, LambdaPP will instead start a ColabFold job to predict the respective structure. As we rely on the current EBI interface to retrieve structures from AlphaFold DB, we implicitly always use the most up to date version provided. At the time of writing, this is AlphaFold DB Release 4 (July 2022) created using the AlphaFold Monomer v2.0 pipeline.

LambdaPP relies on ColabFold 2.1.14 (4) to generate structures using the parameters detailed in listing S9. Implementation details for the backend components can be found here: https://github.com/sacdallago/bio_embeddings/tree/develop/webserver

**Listing S9: Parameters used for structure prediction**

```
use_templates:          False
use_amber:              False
msa_mode:               MMseqs2 (UniRef+Environmental)
num_model:              1
num_recycles:           3
model_order:            [3]
is_complex:             auto
recompile_padding:      1.0
rank_by:                auto
pair_mode:              unpaired+paired
stop_at_score:          85
stop_at_score_below:    40
```
